# Bioinfo-Bench: A Simple Benchmark Framework for LLM Bioinformatics Skills Evaluation

**DOI:** 10.1101/2023.10.18.563023

**Authors:** Qiyuan Chen, Cheng Deng

## Abstract

Large Language Models (LLMs) have garnered significant recognition in the life sciences for their capacity to comprehend and utilize knowledge. The contemporary expectation in diverse industries extends beyond employing LLMs merely as chatbots; instead, there is a growing emphasis on harnessing their potential as adept analysts proficient in dissecting intricate issues within these sectors. The realm of bioinformatics is no exception to this trend. In this paper, we introduce Bioinfo-Bench, a novel yet straightforward benchmark framework suite crafted to assess the academic knowledge and data mining capabilities of foundational models in bioinformatics. Bioinfo-Bench systematically gathered data from three distinct perspectives: knowledge acquisition, knowledge analysis, and knowledge application, facilitating a comprehensive examination of LLMs. Our evaluation encompassed prominent models ChatGPT, Llama, and Galactica. The findings revealed that these LLMs excel in knowledge acquisition, drawing heavily upon their training data for retention. However, their proficiency in addressing practical professional queries and conducting nuanced knowledge inference remains constrained. Given these insights, we are poised to delve deeper into this domain, engaging in further extensive research and discourse. It is pertinent to note that project Bioinfo-Bench is currently in progress, and all associated materials will be made publicly accessible.^1^

## 1 Introduction

Traditional NLP benchmarks have primarily been designed to assess specific and relatively straightforward abilities. However, with the emergence of new capabilities in Large Language Models (LLMs) distinct from smaller models, the focus of evaluation has shifted towards more complex skills, driven by the efforts of researchers. For instance, benchmarks such as MMLU [12], BIG-bench [23], and C-Eval [13] attempt to aggregate a wide range of NLP tasks for holistic assessment. Moreover, various disciplines have constructed their benchmarks tailored to their specific fields through exploration within their respective domains including geoscience [9], radiology [8], and ocean science [5].

Some benchmarks also specifically concentrate on emerging abilities when the models scale, such as inference, mathematical problem-solving [12], and coding [7]. Currently, these benchmarks aim to explore LLMs’ understanding of textual data across various domains including biology and medicine [22, 3] but lack detailed evaluations of domain-specific tasks, limiting the assessment of LLMs’ specific skills. LLMs likely possess a multitude of emerging capabilities that are overshadowed by the singular nature of multiple-choice benchmarks. In this work, we focus on evaluating the advanced abilities of foundational models within the context of bioinformatics.

However, bioinformatics stands out as a promising subfield for integrating biology with LLMs. [27, 6, 19] Numerous studies have explored various aspects not only from the perspective of computer science like medical text analysis [16, **?**], public health data mining [25] and GNN benchmarking [14] but also biological and medical like KVPLM [26], DNA-Pretrain [18], proteome analysis [15, 20, 11], and peptide property prediction [10] within the realm of bioinformatics. Nevertheless, the perspective of LLMs has predominantly focused on handling medical texts, making it challenging for generative artificial intelligence to find suitable entry points and engage effectively. Integrating bioinformatics with LLMs holds immense significance in contemporary scientific research and technological advancements.

Bioinformatics, operating at the crossroads of biology and computational science, is tasked with acquiring, analyzing, and interpreting biological data, particularly in genomics and molecular biology contexts. [4] The emergence of LLMs has revolutionized researchers’ approaches to comprehending biological data, ushering in new avenues for innovation and discovery. Consequently, there is an urgent need for a benchmark that aids in evaluating models’ abilities to analyze and address fundamental issues in bioinformatics. Such a benchmark would enable existing large language models to better assist in tasks like sequence analysis, clinical phenotype prediction, and biological knowledge inference within bioinformatics.

To reduce the gap between LLM development and bioinformatics skills evaluation, we present Bioinfo-Bench, the first simple bioinformatics evaluation suite to thoroughly assess LLMs’ advanced knowledge and problem solving abilities in the bioinformatics. We conduct experiments to evaluate the state-of-the-art LLMs including ChatGPT, Llama, and Galactica on Bioinfo-Bench. In terms of various indicators, LLMs can better complete multiple-choice questions, but they cannot be applied in more in-depth professional tasks. Therefore, we found that in order to truly allow LLMs to penetrate into the practical subject of bioinformatics, more data training in practical application scenarios is needed, rather than simply using the zero-shot abilities and generation capabilities of LLMs.

Finally, in addition to being used as a whole, we also envision Bioinfo-Bench as a set of frameworks. For applied science, we can better evaluate the mastery of LLMs’ knowledge and skills from multiple dimensions, guide developers to understand model capabilities in bioinformatics, and promote LLMs to increasingly help bioinformatics researchers.

**Figure 1:**
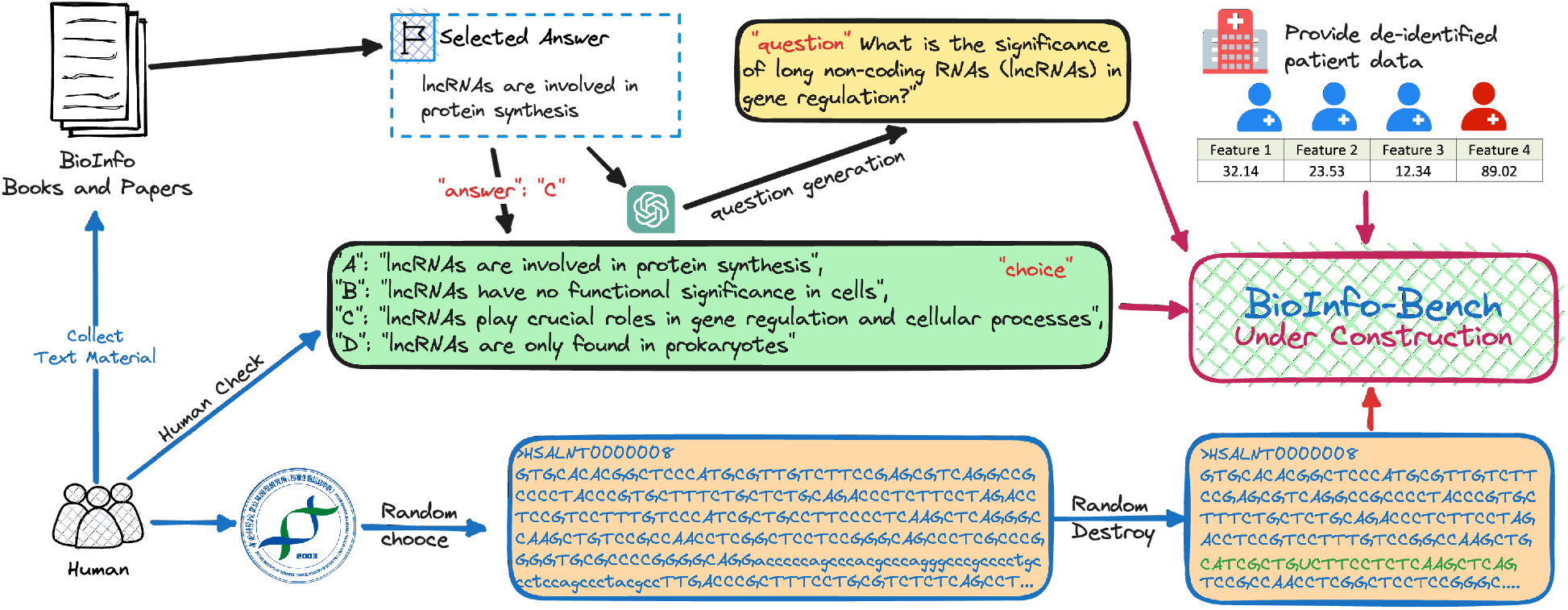
Overview of the Bioinfo-Bench.

## 2 BioInfo-Bench

Assessing various generative models for a specific domain involves subjecting the models to multiple-choice tests. However, interpretations of multiple-choice questions vary widely. Some methods involve concatenating answers into the questions, forming a paragraph, and assessing the perplexity of the generated text. Nevertheless, the phrasing of questions and the combination of options inherently carry biases. Other methods consider the probability distribution of the next token falling on the letters of the options after inputting the question. However, different LLMs also exhibit biases in their understanding of the case sensitivity of options. What’s more, close-source models like ChatGPT, fail to share token probabilities, which we can only use the metric of exactly match, Thus, relying solely on multiple-choice questions to evaluate a model’s understanding of a specific domain is incomplete. Therefore, aligning with bioinformatics needs, we constructed a dataset, BioInfo-Benchmark, comprising 150 multiple-choice questions, 20 sequence verification questions, and 30 disease division tasks.

### 2.1 Data

Initially, we defined various aspects of bioinformatics, outlining ten domains, such as genome analysis, sequence analysis, phylogenetics, structural bioinformatics, gene expression, genetic and population analysis, systems biology, data and text mining, databases and ontologies, bioimage informatics, based on the scope of journals in bioinformatics [1]. We referenced representative bioinformatics tasks, including gene sequence analysis, protein structure prediction, and gene expression data analysis. However, due to the unique nature of the generative models, we could only use generated questions as our evaluation criteria. For multiple-choice questions, we gathered authoritative bioinformatics literature from the internet. We used these sources to formulate questions for ChatGPT, with the constraint that the answers must be found within the original texts. We collected 100 high-quality bioinformatics papers for each domain, creating 15 question seeds. ChatGPT produced three similar answers in generating responses, forming four candidate responses. Finally, we curated a pool of multiple-choice questions through manual selection, accumulating 150 questions in ten bioinformatics subdomains. We name this part of benchmarks as Bioinfo-Bench-qa.

For sequence verification questions, we collect 20 RNA sequences from LncBook [17] made by NGDC, sharing an example. For each sequence, we randomly re-order, delete, add, and replace the RNA base pairs tokens to ask the LLMs for advice on which action has proceeded. This task mainly examines the model’s ability to utilize sequence memory and common sense of bioinformatics. We name this part of benchmarks as Bioinfo-Bench-seq.

**Figure 2:**
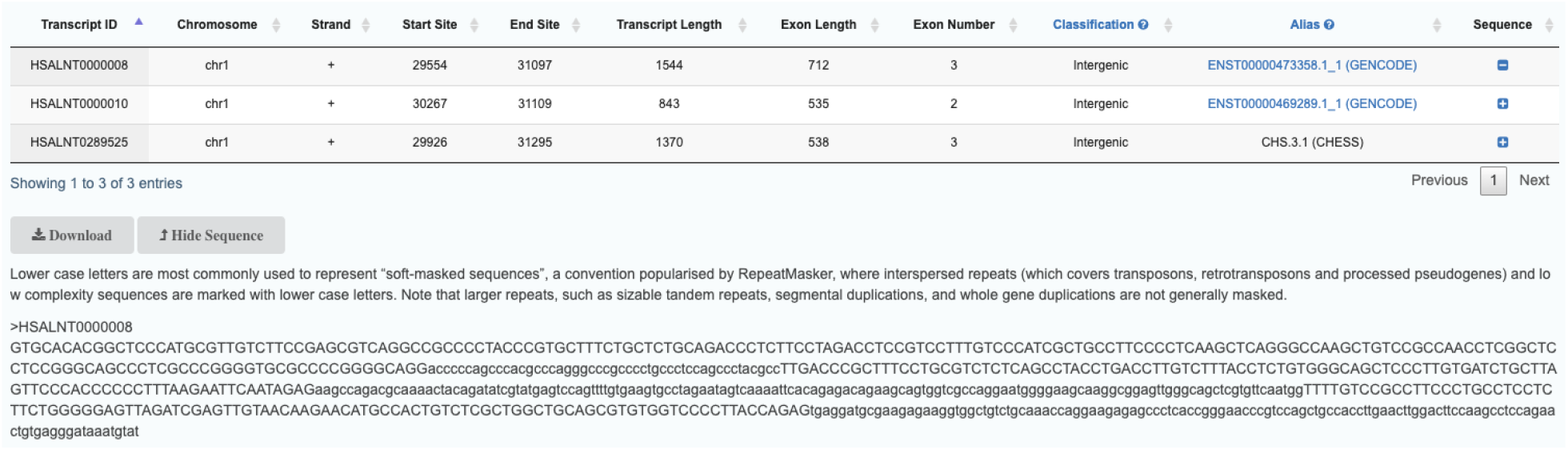
Introduction to the source of task Bioinfo-Bench-seq.

We collect 30 patient de-identified data from a cooperating hospital for disease division tasks. Each patient is characterized by four attributes, including biological targets (a detailed introduction will be added later as this target involves another unpublished paper). The LLM receives five positive and five negative signals as supervisory signals. This setup enables the model to analyze new patient data and determine the patient’s health status as a few-shot learning. The primary objective of this task is to assess the practical data analysis capabilities of the LLM. We name this part of benchmarks as Bioinfo-Bench-div.

### 2.2 Metric

To ensure a fair and impartial evaluation of the models, we applied consistent treatment to perplexity and the probability of the next token. Specifically, we selected the optimal score between the two metrics.

As for perplexity (ppl), ppl is one of the most common metrics for evaluating language models. This type of metric applies specifically to autoregressive models or decoder only model and is not well defined for encoder-only model e.g. BERT. ppl is defined as the exponentiated average negative log-likelihood of a sequence. If we have a tokenized sequence, then the perplexity is,

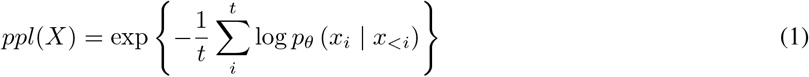

Therefore, for a multiple-choice question, we splice the options back together to calculate the degree of confusion of the option vocabulary under the generated representation of the model in the sentence obtained after splicing.

However, according to the generative language model, intuitively, it can be thought of as an assessment of a model’s ability to predict uniformly across a specified set of tokens in the corpus. Importantly, this means that the tokenization process has a direct impact on the complexity of the model. We take this into consideration when comparing different models, so we added the probability of the next token as another evaluation metric for multiple-choice questions.

Evaluating the model output is also equivalent to the exponentiation of the cross-entropy between the data and model predictions. The model takes the prefix (in our evaluation context, the question) as input for prediction. The model generates a vector equal to the vocabulary, containing probabilities assigned to each word. Subsequently, we query the respective token IDs using the tokenizer of each model and compare the probabilities to determine the optimal probability. At the same time, for ChatGPT, this probability is not convenient to output. Therefore, we only use the exact match form to evaluate ChatGPT.

## 3 BioInfo-Bench

### 3.1 Baseline

There is a large quantity of large language models in the world.

- **ChatGPT**: We use the API of the 2023 March version of ChatGPT (gpt-3.5-turbo-0301). ChatGPT represents a multitude of non-open-source language models where users are limited to receiving responses without the ability to access the underlying code. In many cases, such language models require scrutiny through trust and safety (TnS) checks to ensure reliability and legality, imposing certain constraints on their inherent capabilities.
- **Llama-7B**: Llama 7B is one of the foundation language models developed by Meta AI. It is part of the Llama 2 collection, which includes models ranging from 7B to 70B parameters. Llama represents a group of outstanding open-source models, allowing users to conduct further fine-tuning to explore their specific domains. Such language models exhibit strong adaptability. We opted for the 7b release as it marks the inception of emerging phenomena and serves as a consumable language model accessible for laboratory-level scientific explorations.
- **Galactica-30B**: Galactica-30B, created by the Papers with Code team at Meta AI, is a powerful scientific LLM. It excels in citation prediction, scientific QA, mathematical reasoning, summarization, document generation, molecular property prediction, and entity extraction. Galactica, developed through specialized text training, offers the academic community a distinctive perspective on knowledge, reshaping the distribution of scholarly information. While this model demonstrates a high level of specialization, its expressive capabilities may not necessarily reach the forefront of the field.

### 3.2 Analysis

As mentioned above, we use the baselines and metrics to evaluate ChatGPT, Llama, and Galactica. The results are shown in Table 1.

**Table 1:**
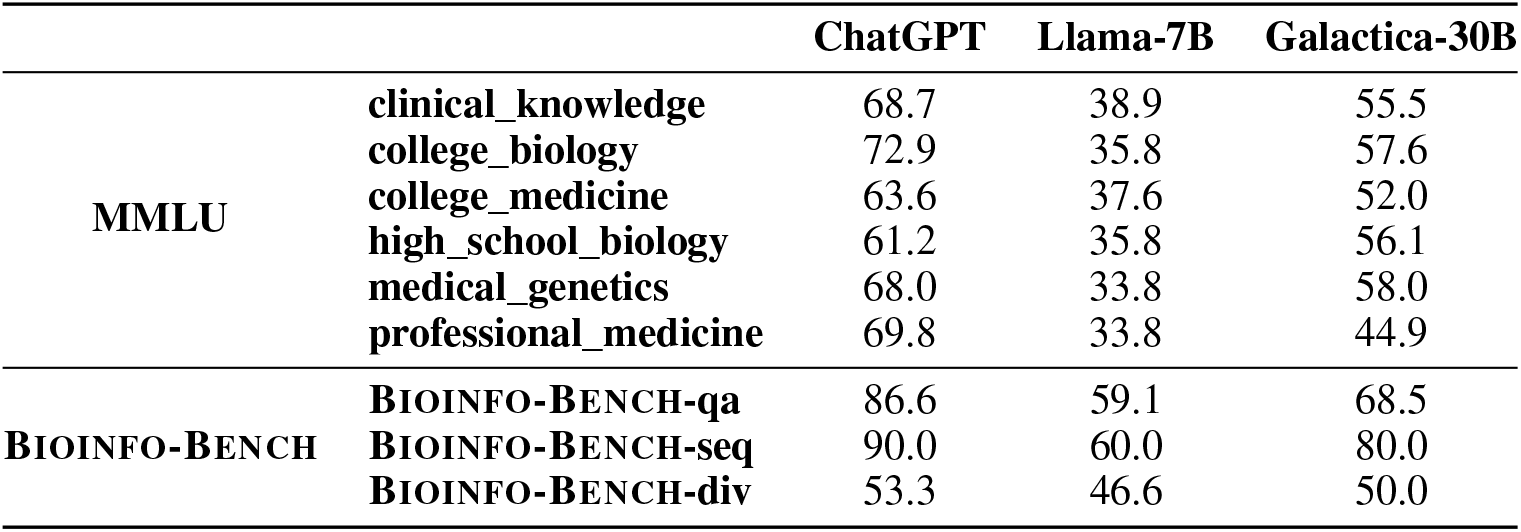
The result of baseline models on biology-related and medicine-related MMLU and BioInfo-Bench.

The survey results indicate that these Large Language Models (LLMs) exhibit outstanding performance in knowledge acquisition, primarily relying on their training data to retain information. However, their ability to solve real-world professional problems and engage in nuanced knowledge inference remains limited. It is worth noting that the scale of the current benchmark needs further expansion. Additionally, the complexity and difficulty of the selected questions pose challenges. In the future, we will conduct an in-depth analysis of question selection to achieve optimal evaluation outcomes.

## 4 Discussion

### 4.1 Data collection

One of the key aspects where the cooperation between bioinformatics and LLMs is crucial lies in the analysis of vast biological datasets. With the continuous advancements in high-throughput technologies, biological data is being generated at an unprecedented rate. LLMs, equipped with their natural language processing capabilities and machine learning algorithms, have the potential to handle and interpret this immense volume of data efficiently. They can aid in tasks such as sequence analysis, protein structure prediction, and functional genomics, enabling researchers to extract meaningful insights from complex biological information.

### 4.2 Cooperation

Furthermore, integrating LLMs in bioinformatics facilitates effective communication and collaboration between experts from diverse fields. By enabling researchers to articulate complex biological concepts in a language understandable to computational scientists and vice versa, LLMs bridge the gap between domain-specific knowledge and computational methodologies. This interdisciplinary collaboration fosters innovation and accelerates the pace of scientific discoveries in both bioinformatics and related computational disciplines.

In addition, LLMs enhance the accessibility of biological knowledge to a broader audience, including researchers, clinicians, educators, and policymakers. By comprehensibly processing and presenting biological information, these models democratize access to critical scientific insights, promoting a more informed decision-making process in various sectors, such as healthcare, biotechnology, and environmental conservation.

In summary, integrating bioinformatics with large language models improves analytical capabilities and promotes interdisciplinary collaboration, knowledge dissemination, and informed decision-making. This cooperation represents a paradigm shift in how biological data is analyzed, interpreted, and applied, paving the way for transformative advancements in biology and computer science.

### 4.3 Future of Bioinfo-Bench

We plan to approach tasks related to bioinformatics demands in the future systematically. We aim to create an evaluation framework based on Bioinfo-Bench primarily centered around practical tasks, with multiple-choice questions as supplementary elements. From a practical perspective, we will allow models to utilize additional tools, such as Code Integrator [2, 24] and Toolformer [21]. Furthermore, we will continuously enrich our question collection. Given the increasing prevalence of foundational bioinformatics courses, including those at the undergraduate level in various universities, we intend to enhance our evaluation framework from an educational perspective.

At the appropriate juncture, we will introduce evaluation benchmarks specifically tailored for vision modalities. This initiative aims to comprehensively assess the abilities of LLMs with visional perception capabilities in bioinformatics. Lastly, we are committed to the ongoing maintenance of Bioinfo-Bench project, supporting the convergence of bioinformatics and LLMs in this specialized domain.

## 5 Conclusion

In this article, we introduce a novel and straightforward framework to assess the capabilities of LLMs in bioinformatics. The aim is to examine whether these LLMs can effectively assist experts in bioinformatics. We constructed 200 questions covering multiple-choice, sequence verification, and analytical problem-solving tasks utilizing machine learning techniques specific to bioinformatics. Experimental results indicate that LLMs demonstrate proficiency in answering questions by relying on their memorization abilities derived from pre-training corpora. However, they exhibit notable limitations in tackling questions requiring specialized reasoning and analysis. Consequently, for such nuanced issues, we plan to conduct further analysis and delve deeper into exploration while continually updating our project. Our initiative will remain open-source on GitHub, ensuring accessibility and transparency for the academic community.

## Footnotes

1 https://github.com/cinnnna/bioinfo-bench

